# NaWRKY3 is a master transcriptional regulator of defense network against *Alternaria alternata* in *Nicotiana attenuata*

**DOI:** 10.1101/2022.11.29.518390

**Authors:** Zhen Xu, Shuting Zhang, Jinsong Wu

**Affiliations:** Yunnan Key Laboratory for Wild Plant Resources, Kunming Institute of Botany, Chinese Academy of Sciences, Kunming, 650201, China; University of Chinese Academy of Science, Beijing 10049, China; Yunnan Key Laboratory for Fungal Diversity and Green Development, Kunming Institute of Botany, Chinese Academy of Sciences, Kunming, 650201, China

**Keywords:** Alternaria, *BBL*, ethylene, *F6’H1*, HGL-DTGs, jasmonate, lncRNA, plant resistance, phytoalexin, scopoletin, *Rboh*

## Abstract

WRKY transcription factors are involved in plant defense responses against pathogens. However, no WRKYs have been reported yet in resistance of *Nicotiana* species to *Alternaria alternata*, a necrotrophic fungal pathogen causing brown spot disease. Here, we found that silencing *NaWRKY3* lead to wild tobacco *Nicotiana attenuata* highly susceptible to *A. alternata*. Combination of transcriptome, electrophoretic mobility shift, ChIP-qPCR and dual-LUC analyses, we uncovered that NaWRKY3 bound to many defense genes’ promoter and activated their expression. Target genes included: 1) *lipoxygenases 3*, *ACC synthase 1* and *ACC oxidase 1*, three key enzyme genes for JA and ethylene biosynthesis which were critical for *A. alternata* resistance; 2) *feruloyl-CoA 6’-hydroxylase 1* (*NaF6’H1*), the key enzyme gene for phytoalexins against *A. alternata*, scopoletin and scopolin; and 3) three *A. alternata* resistance genes, long non-coding RNA (*LncRNA L2*), *NADPH oxidase* (*NaRboh D*) and *berberine bridge-like* (*NaBBL28*). Silencing *LncRNA L2* reduced *A. alternata*-induced levels of JA and *NaF6’H1* expression. *NaRboh D*-silenced plants were strongly impaired in ROS production and stomata closure responses. *NaBBL28* was the first *A. alternata* resistance BBLs identified and was involved in HGL-DTGs hydroxylation. Finally, NaWRKY3 could bind to its own promoter but acted as a transcriptional repressor. Thus we demonstrated that NaWRKY3 is a fine-tuned master regulator of defense network against *A. alternata* in *N. attenuata* by regulating different signaling pathways and defense metabolites. For the first time, such an important WRKY was identified in *Nicotiana* species, providing new insight into defense mechanism of *Nicotiana* plants to *A. alternata*.

## Introduction

To cope with pathogens, different layers of plant defense response are activated, including pathogen recognition, activation of phytohormone and reactive oxygen species (ROS) signaling pathways, production of antimicrobial metabolites and proteins, stomata closure, and physical reinforcement of cell walls through production of callose and lignin (Camejo et al., 2016; Zhang et al., 2018).

Phytohormones and Rboh-based ROS are important signaling molecules, playing pivotal roles in regulating plant defense response. The jasmonate (JA) and ethylene signaling pathways are usually associated with plant defense against necrotrophic pathogens (Glazebrook, 2005). *Alternaria alternata* (tobacco pathotype) is a necrotrophic fungal pathogen causing brown spot diseases in *Nicotiana* species (LaMondia, 2001). In wild tobacco *Nicotiana attenuata*, lipoxygenases 3 *(NaLOX3*), allene oxide synthase (*NaAOS*) and allene oxide cyclase (*NaAOC*) are three key enzyme genes of JA biosynthesis (Halitschke and Baldwin, 2003; Kallenbach et al., 2012; Sun et al., 2014). *N. attenuata* plants increased their JA levels in response to *A. alternata* infection at 1 dpi, and silencing *AOC* or receptor gene *COI1* lead to plants highly susceptible to the fungus (Sun et al., 2014). Ethylene is produced from S-adenosyl-L-methionine (SAM) with the help of ACC synthase (ACS) and ACC oxidase (ACO). Both NaACS and NaACO are encoded by multi-gene families in *N. attenuata* (von Dahl et al., 2007). *NaACSs* and *NaACOs* are strongly up-regulated after *A. alternata* infection, and bigger lesions are developed in *ACOs*-silenced plants, suggesting that ethylene signaling pathway plays an essential role in the resistance of *N. attenuata* to *A. alternata* (Sun et al., 2017). Reactive oxygen species (ROS) act as important signaling molecules during pathogen attack (Camejo et al., 2016). Rboh, an integral membrane protein, is the homologue of NADPH oxidase generating O_2_^−^ during pathogen attack. NbRboh A and NbRboh B are essential for ROS production against oomycete pathogen *Phytophthora infestans* (Adachi et al., 2015). GbRboh5/18 enhances cottons’ resistance against *Verticillium dahliae* by elevating the levels of ROS (Chang et al., 2020). In *N. attenuata*, NaRboh D is the major ROS source after herbivore attack, and confers plant resistance to *Spodoptora littoralis* (Wu et al., 2013). However, whether this NaRboh D is involved in *A. alternata* resistance is unknown.

Phytoalexins, as direct chemical weapons, are produced by plants *de novo* in response to pathogen attack. Camalexin is the most prominent phytoalexin in *Brassicaceae* (He et al., 2019; Wu, 2020; Zhou et al., 2022). Capsidiol is an important phytoalexin against *A. alternata* in *N. attenuata* (Song et al., 2019). Scopoletin, known as anti-carcinogenic and anti-viral natural coumarin, is demonstrated as a phytoalexin against *A. alternata* regulated by JA and ethylene signaling pathways (Sun et al., 2014; Sun et al., 2017).

WRKYs are the biggest and most important transcription factor family in plants involved in plant immunity. They are characterized by the conserved WRKYGQK and zinc finger motifs, recognizing *cis*-elements W-boxes in the promoter of target genes and activating or inhibiting their expression (Wani et al., 2021). AtWRKY33 is required for resistance to necrotrophic fungal pathogen *Botrytis cinerea* via regulating phytohormones and camalexin (Birkenbihl et al., 2012). AtWRKY57 is a repressor of *B. cinerea* resistance due to activating JA signaling pathway repressors JAZ1 and JAZ5 (Jiang and Yu, 2016). AtWRKY46, AtWRKY70, and AtWRKY53 positively regulate basal resistance to *Pseudomonas syringae*, while AtWRKY8 represses resistance to this pathogen (Hu et al., 2012). However, to date no WRKYs have been reported to play a role in the defense response of *Nicotiana* species to *A. alternata*.

Currently, it is still unclear how phytohormone and ROS signaling pathways, and phytoalexins are coordinately regulated in *Nicotiana* species during *A. alternata* attack. In this study, we identified the transcription factor NaWRKY3 as a master transcriptional regulator of defense network against *A. alternata* in *N. attenuata*. NaWRKY3 enhanced the expression of target genes involved in different layers of plant defense, including those for directly defense response like phytoalexins, and immune signals like phytohormones and ROS from respiratory burst oxidase, *A. alternata* resistance LncRNA *L2* and *NaBBL28*, while repressed its own transcription. For the first time, we identified such an important WRKY involved in so many defense responses in *N. attenuata*, thus providing new insight into defense mechanism of *Nicotiana* species to *A. alternata*.

## Results

### NaWRKY3 is required for *A. alternata* resistance in *N. attenuata*

Previously, NaWRKY3 was reported to be involved in herbivore resistance in *N. attenuata* (Skibbe et al., 2008). We also found that NaWRKY3 transcripts were significantly up-regulated in *N. attenuata* after *A. alternata* inoculation during transcriptome analysis (Song et al., 2019). This result was also confirmed by real time PCR that it was expressed at low levels in healthy, uninfected source-sink transition leaves (0 leaves) of wild-type (WT) plants; but increased significantly at 1 day post inoculation (dpi) and reached around 10-fold at 3 dpi (Fig. 1a).

**Fig. 1.**
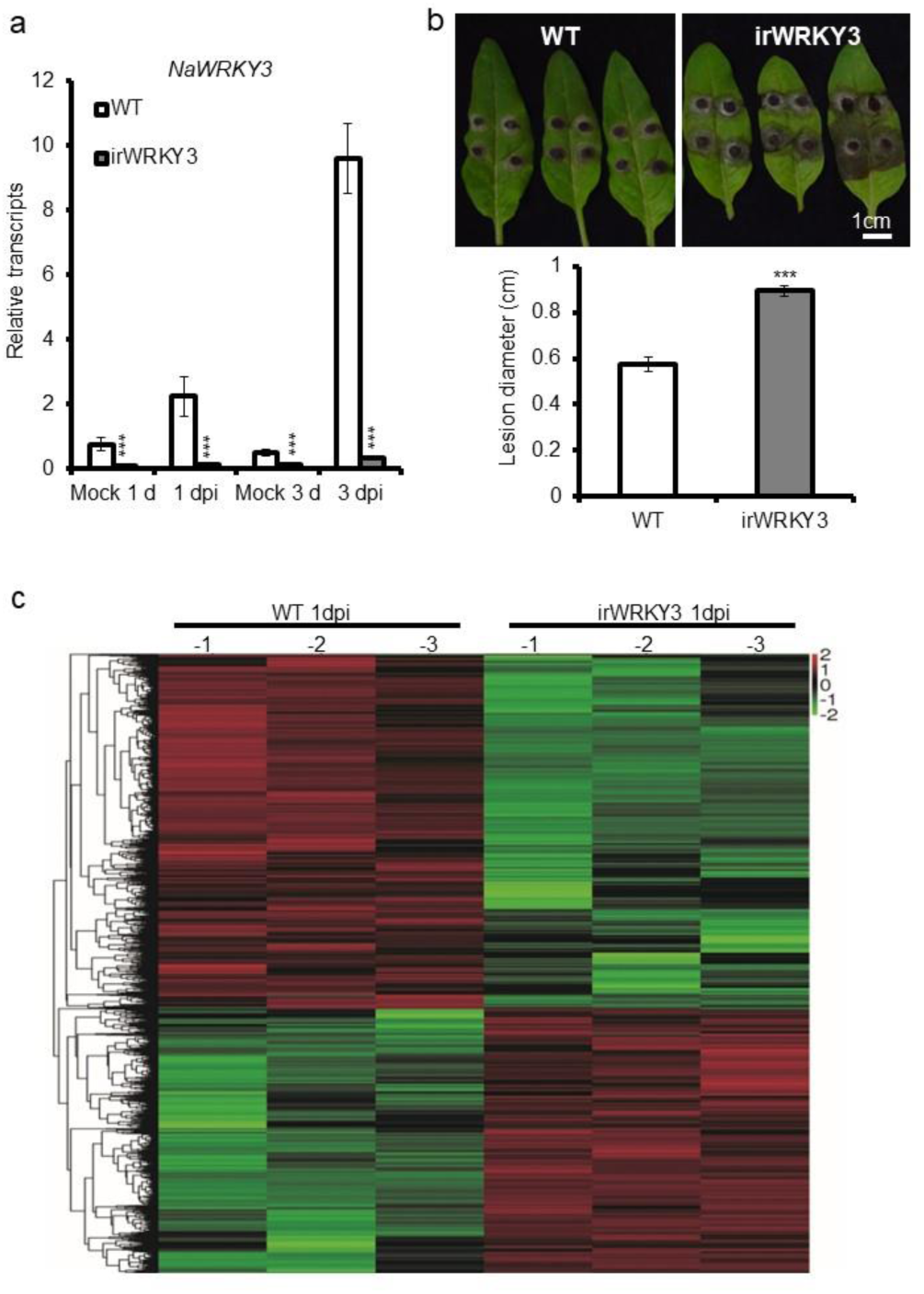
NaWRKY3 is required for *A. alternata* resistance in *N. attenuata*. **(a)** Mean (±SE) relative *A. alternata*-induced *NaWRKY3* transcripts as measured by qPCR in five replicates of source-sink transition leaves (0 leaves) at 1 and 3 dpi. Asterisks indicated levels of significant difference between WT and irWRKY3 plants with the same treatments (Student’s *t*-test: ***P<0.005) **(b)** Photographs of infected WT and irWRKY3 leaves were taken at 4 dpi (upper panel); Mean (±SE) diameter of necrotic lesions of 0 leaves of WT and irWRKY3 plants at 4 dpi (n = 20; lower panel). Asterisks indicated the levels of significant differences between WT and irWRKY3 plants with the same treatments (Student’s *t*-test: ***, p < 0.005). **(c)** Heatmap of DEGs in three replicates of 0 leaves of WT and irWRKY3 plants after *A. alternata* infected at 1 dpi.

To determine the function of NaWRKY3 in *A. alternata* resistance, stable transgenic plants silenced with *NaWRKY3* (irWRKY3), which were generated previously by Skibbe et al. (2008), were used for *A. alternata* inoculation. In these plants, *NaWRKY3* transcripts were decreased by around 87% in mock control, and 96% at 3 dpi (Fig. 1a). As expected, the 0 leaves of irWRKY3 plants showed bigger lesions than those of WT at 4 dpi (Fig. 1b), which marked increased susceptibility to *A. alternata*. Thus, our results show that NaWRKY3 plays an important role in defense against *A. alternata*.

### Transcriptome analysis of irWRKY3 plants after *A. alternata* inoculation

We performed a transcriptome analysis through RNA sequencing to obtain a deep view of the transcriptional reprogramming mediated by NaWRKY3 upon *A. alternata* challenge. A total of 12 samples were collected from 0 leaves of WT and irWRKY3 plants with or without *A. alternata* at 1 dpi, respectively.

Silencing *NaWRKY3* resulted a massive transcriptional reprogramming. *A. alternata* infected 0 leaves of irWRKY3 plants up-regulated 3669 genes and repressed 4921 genes at 1 dpi when compared to WT plants with the same treatments (Fig. 1c). We especially interested in genes related to defense that were down-regulated in irWRKY3 plants, including genes involved in JA, ethylene, ROS, long non-coding RNA, phytoalexins and berberine bridge-like. We thus selected them for further analysis as below.

### NaWRKY3 is required for JA accumulation and the expression of JA synthesis gene *NaLOX3* after *A. alternata* inoculation

The JA levels were increased form 0.55 ng/g to 70 ng/g fresh weight in 0 leaves of WT plants after 1 dpi, but *A. alternata*-induced JA levels in irWRKY3 plants were only 58% of those of WT plants with the same treatments (Fig. 2a). Similarly, *A. alternata*-induced JA-Ile was also significantly attenuated in irWRKY3 plants (Fig. 2a).

**Fig. 2.**
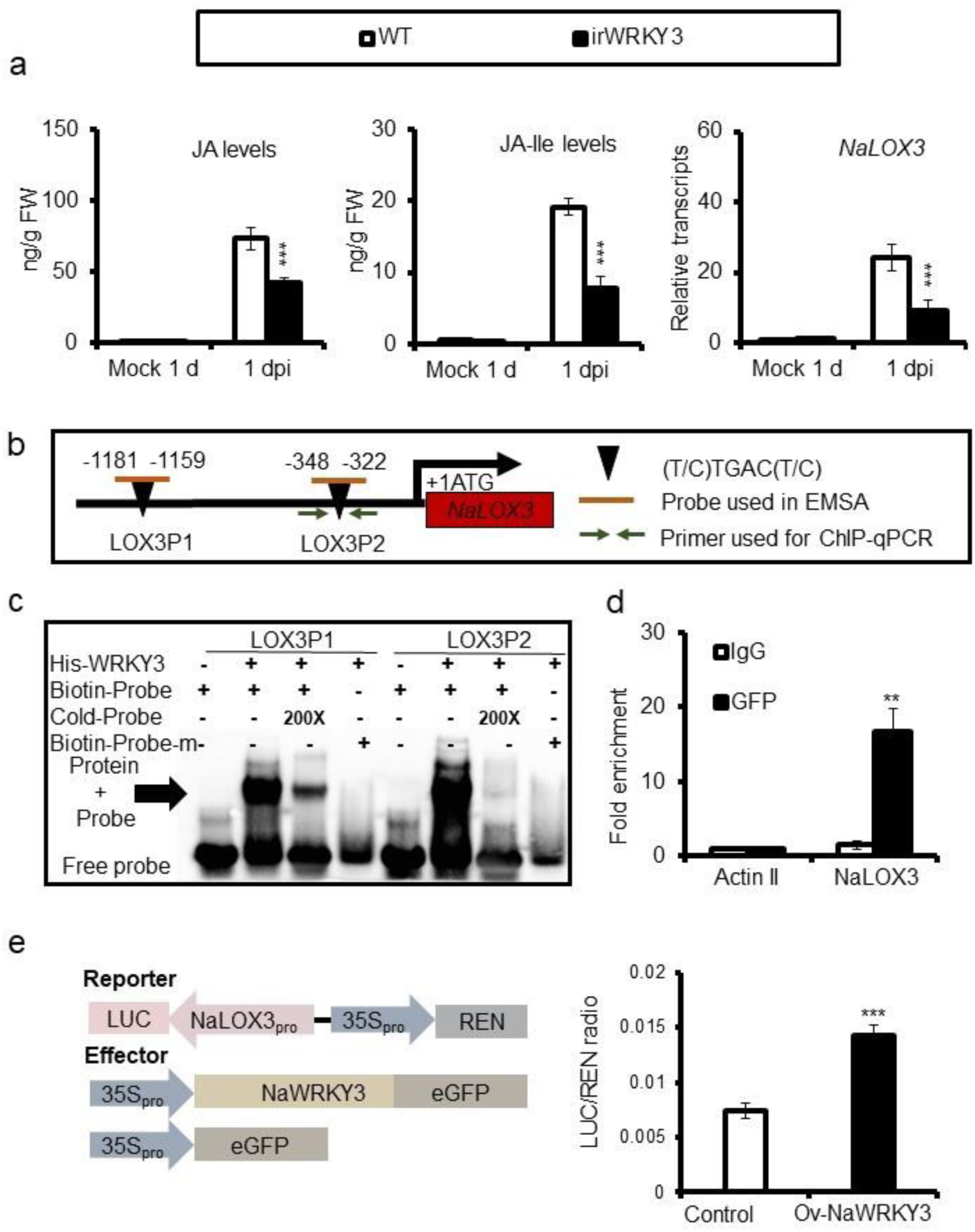
NaWRKY3 is required for JA accumulation and the expression of *NaLOX3* after *A. alternata* inoculation. **(a)** Mean (±SE) relative *A. alternata*-induced JA/JA-Ile levels and *NaLOX3* transcripts in five replicates of 0 leaves of WT and irWRKY3 plants at 1 dpi. JA and JA-Ile levels were determined by LC-MS/MS, NaLOX3 transcripts were measured by qPCR. Asterisks indicated the levels of significant differences between WT and irWRKY3 plants with the same treatments (Student’s *t*-test: ***P<0.005). **(b)** Schematic diagram of the promoter of *NaLOX3*. Black triangles indicated the W-box motifs. Short orange lines indicated DNA probes used for EMSA, and green arrows indicated primers used for ChIP-qPCR assays. The translational start site (ATG) was shown at position +1. **(c)** EMSA showed that His-NaWRKY3 bind to the promoter of *NaLOX3* directly. Labeled probes incubated without His-NaWRKY3 served as a negative control. 200-fold excess of unlabeled probes were used for competition. Biotin-Probe-m was the probe with mutated W-box motif. **(d)** Mean (±SE) fold enrichment on the NaWRKY3-eGFP plants in ChIP-qPCR. DNA from NaWRKY3-eGFP over-expressed plants at 1 dpi was immune-precipitated by GFP or IgG antibody, and was further analyzed by qPCR with indicated primers. The immune-precipitated DNA from IgG antibody was served as control, and asterisks indicated levels of significant difference between negative controls and experimental group (Student’s *t*-test: **P<0.01). **(e)** Schematic diagram of constructs for effectors and reporters (left panel), and mean (±SE) relative NaLOX3_pro_::LUC transient transcription activity activated by NaWRKY3 in *N. benthamiana* plants (right panel). The *Agrobacterium* with NaLOX3_pro_::LUC reporter was co-injected with those with indicated effector into leaves of *N. benthamiana*. After 36 h, values represented the NaLOX3_pro_::LUC activity relative to the internal control (35S::REN activity) were obtained. Asterisks indicated significant differences between control and Ov-NaWRKY3 samples (Student’s *t*-test; n=4; ***P<0.005).

Lipoxygenase 3 (NaLOX3) was an important enzyme in JA biosynthesis (Halitschke and Baldwin, 2003). Transcriptome data indicated *NaLOX3* was decreased in irWRKY3 plants at 1 dpi. Indeed, real-time PCR revealed that levels of *A. alternata*-induced *NaLOX3* in 0 leaves of irWRKY3 plants were only 38% of that of WT plants (Fig. 2a), indicating that NaWRKY3 was required for the fungus-induced expression of *NaLOX3*. Meanwhile, another enzyme gene *NaAOS* for JA biosynthesis did not alter its expression in irWRKY3 plants (Supplemental Fig. 1a).

Biotin-labeled probes were synthesized according to W-boxes in the promoter of *NaLOX3* (Fig. 2b). Recombinant NaWRKY3 fused with 6x His was purified for electrophoretic mobility shift assay (EMSA). Our results indicated that NaWRKY3 could specifically bind to two selected probes (LOX3P1 and LOX3P2), but not to their mutated ones (Biotin-Probe-m). However, these bindings were greatly impaired when 200 times of unlabeled cold probes were applied (Fig. 2c). These results indicated that NaWRKY3 could specifically bind to *NaLOX3* promoter directly *in vitro*.

To further confirm the binding of NaWRKY3 and *NaLOX3* promoter *in vivo*, *NaWRKY3-eGFP* over-expressed transgenic plants were used for ChIP-qPCR. Primers were designed according to the binding sites of NaWRKY3 obtained from EMSA results (Fig. 2b and 2c). Our results showed that DNA regions which were immuno-precipitated by anti-GFP antibody, were more enriched around the W-boxes of *NaLOX3* promoter than those of Actin II (Fig. 2d), suggesting that NaWRKY3 could directly bind to the promoter of *NaLOX3 in vivo*.

A dual-LUC reporter assay was employed to investigate whether NaWRKY3 activated the expression of *NaLOX3* promoter. The expression of *LUC* (reporter) was driven by the promoter of *NaLOX3* (NaLOX3_pro_), and *REN* driven by 35S promoter acted as an internal control (Fig. 2e). When *NaWRKY3* was over-expressed under the control of 35S promoter (effector; 35S::NaWRKY3), a significant increasing in LUC/REN ratio was observed at 36 h, indicating that NaWRKY3 could activate *NaLOX3* promoter (Fig. 2e).

Taken together, our results indicate that NaWRKY3 is a transcriptional activator of JA synthetic gene *NaLOX3* and regulates the expression of *NaLOX3* by directly binding to W-boxes within its promoter.

### NaWRKY3 regulates an *A. alternata* resistance lncRNA *L2*

Several long non-coding RNAs (lncRNAs) were found to be dramatically up-regulated in WT plants at 1 dpi, but were not in irWRKY3 plants during transcriptome analysis. qPCR results confirmed that XR_002068323.1 (*L2*), XR_002067418.1 (*L6*) and XR_002066427.1 (*L4*) were strongly induced in 0 leaves of WT plants after *A. alternata* inoculation at 1 and 3 dpi, but these inductions were greatly reduced in irWRKY3 plants at both time points (Supplemental Fig. 2 and Fig. 3c). We thus silenced these three lncRNAs via VIGS separately (Supplemental Fig. 3a). Our results revealed that *L2* was required for *N. attenuata* resistance to *A. alternata*, as bigger lesions were developed in *L2*-silenced plants (Supplemental Fig. 3b). Thus, *L2* was chosen for further analysis.

To further explore the role of *L2*, stable transformed RNAi (RNA interference) plants were generated. We obtained two T2 transgenic *L2* RNAi lines, irL2-1 and irL2-3. qPCR showed *L2* expression were significantly decreased in both RNAi lines at 1 dpi (Fig. 3a). As expected, the enhanced susceptibility to *A. alternata* was observed in irL2-1 and irL2-3 plants (Fig. 3b). Interestingly, *A. alternata*-induced *NaF6’H1* expression, JA and JA-Ile levels were decreased in irL2-1 and irL2-3 plants (Fig. 3a). These results indicated that silencing *L2* impaired *A. alternata*-induced JA and *NaF6’H1* expression.

**Fig. 3.**
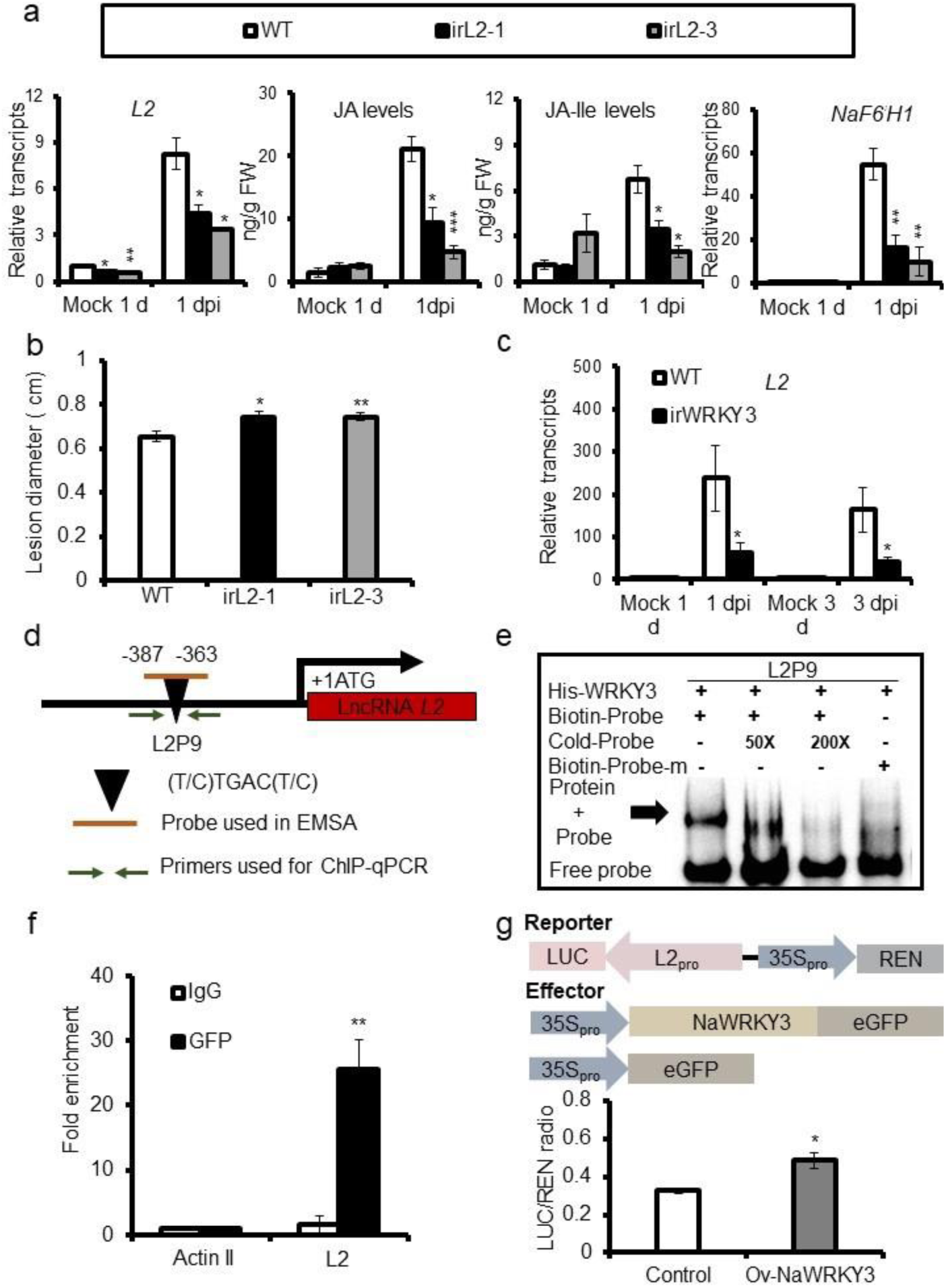
NaWRKY3 regulates an *A. alternata* resistance lncRNA *L2.* **(a)** Mean (±SE) relative *A. alternata*-induced transcripts of lncRNA *L2* and *NaF6’H1*, and JA and JA-Ile levels as measured in five replicate 0 leaves of WT and two lncRNA *L2* RNAi lines at 1 dpi. Asterisks indicated levels of significant difference between WT and lncRNA *L2* RNAi lines (Student’s *t*-test: *P<0.05, **P<0.01, ***P<0.005). **(b)** Mean (±SE) diameter of necrotic lesions of 20 biological replicate 0 leaves of WT and irL2 (irL2-1 and irL2-3) plants at 4 dpi (n = 20). Asterisks indicated the levels of significant differences between WT and lncRNA *L2* RNAi plants with the same treatments (Student’s *t*-test: *P<0.05, **P<0.01). **(c)** Mean (±SE) relative *A. alternata*-induced lncRNA *L2* transcripts as measured by qPCR in five replicate leaves of WT and irWRKY3 plants at 1 and 3 dpi. Asterisks indicated the levels of significant differences between WT and irWRKY3 plants with the same treatments (Student’s *t*-test: *P<0.05). **(d)** Schematic diagram of the promoter of lncRNA *L2*. Black triangle indicated the W-box motif. Short orange line indicated DNA probe used for EMSA, and green arrows indicated primers used for ChIP-qPCR assays. The translational start site (ATG) was shown at position +1. **(e)** EMSA showed that His-NaWRKY3 could bind to the promoter of lncRNA *L2* directly. 50- and 200-fold excess of unlabeled probes were used for competition. Biotin-Probe-m was the probe with mutated W-box motif. **(f)** Mean (±SE) fold enrichment on the NaWRKY3-eGFP plants in ChIP-qPCR. DNA from *NaWRKY3-eGFP* over-expressed plants at 1 dpi was immune-precipitated by GFP or IgG antibody, and was further analyzed by real time PCR with indicated primers. The immune-precipitated DNA from IgG antibody was served as control, and asterisks indicated levels of significant difference between negative controls and experimental group (Student’s t-test: **P<0.01). **(g)** Schematic diagram of constructs for effectors and reporters (left panel), and mean (±SE) relative L2_pro_::LUC transient transcription activity activated by NaWRKY3 in *N. benthamiana* plants (right panel). The *Agrobacterium* with L2_pro_::LUC reporter was co-injected with those with indicated effector into leaves of *N. benthamiana*. After 36 h, values represented the L2pro::LUC activity relative to the internal control (35S::REN activity) were obtained. Asterisks indicated significant differences between control and Ov-NaWRKY3 samples (Student’s *t*-test; n=4; *P<0.05).

Next, we performed EMSA and ChIP-qPCR to investigate the regulation of *L2* by NaWRKY3. EMSA results indicated that NaWRKY3 could specifically bind to the probe L2P9 which was designed from *L2* promoter containing a W-box, as it could not bind to the mutated L2P9 (Biotin-Probe-m; Fig. 3d and 3e), and the binding of NaWRKY3 to L2P9 was attenuated when 50 times of cold unlabeled probes were additionally added and abolished when 200 times of cold probes were applied (Fig. 3e). Further ChIP-qPCR assay using anti-GFP antibody and *NaWRKY3-eGFP* over-expressed transgenic plants indicated that NaWRKY3 could bind to the promoter of *L2 in vivo* (Fig. 3f).

The activation of *L2* promoter by NaWRKY3 was also tested by dual-LUC reporter assay. A significant increasing in LUC/REN ratio was observed when NaWRKY3 was over-expressed for 36 h (Fig. 3g), suggesting that NaWRKY3 could activate the promoter of *L2*.

Thus, we identify lncRNA *L2*, a direct target gene of NaWRKY3. We demonstrate that this lncRNA is required for *A. alternata* resistance likely by affecting JA and scopoletin responses.

### Silencing NaWRKY3 impairs expressions of ethylene biosynthetic genes and ethylene emission during *A. alternata* inoculation

After *A. alternata* inoculation, 0 leaves of *N. attenuata* emitted around 12 nL/g FW ethylene in 24 h (Fig. 4a). However, *A. alternata*-elicited ethylene production was reduced by 70% in irWRKY3 plants (Fig. 4a), suggesting NaWRKY3 was involved in this fungus-induced ethylene signaling.

**Fig. 4.**
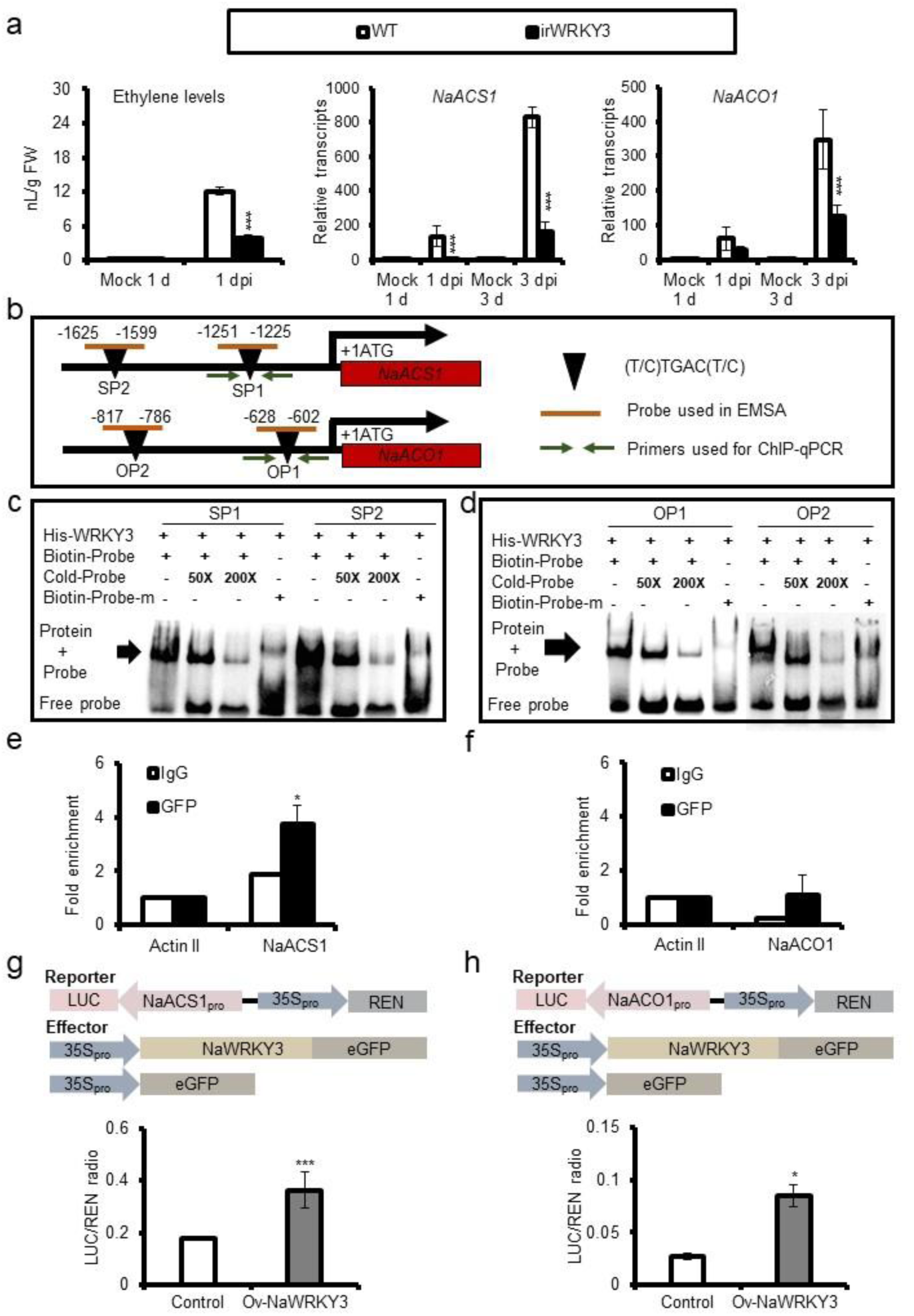
NaWRKY3 is required for ethylene emission and the expression of *NaACS1* and *NaACO1* after *A. alternata* inoculation. **(a)** Mean (±SE) relative *A. alternata*-induced ethylene production, and *NaACS1*/*NaACO1* transcripts in five replicate 0 leaves of WT and irWRKY3 plants. Ethylene levels were determined by GC-MS/MS, *NaACS1* and *NaACO1* transcripts as measured by qPCR. Asterisks indicated the levels of significant differences between WT and irWRKY3 plants with the same treatments (Student’s *t*-test: ***P<0.005). **(b)** Schematic diagram of the promoter of *NaACS1* and *NaACO1*. Black triangle indicated the W-box motif. Short orange lines indicated DNA probes used for EMSA, and green arrows indicated primers used for ChIP-qPCR assays. The translational start site (ATG) was shown at position +1. **(c)** and **(d)** EMSA showed that His-NaWRKY3 could bind to the promoter of *NaACS1* and *NaACO1* directly. 50- and 200-fold excess of unlabeled probes were used for competition. Biotin-Probe-m was the probe with mutated W-box motif. **(e)** and **(f)** Mean (±SE) fold enrichment on the NaWRKY3-eGFP plants in ChIP-qPCR. DNA from *NaWRKY3-eGFP* over-expressed plants at 1 dpi was immune-precipitated by GFP or IgG antibody, and was further analyzed by real time PCR with indicated primers. The immune-precipitated DNA from IgG antibody was served as control, and asterisks indicated levels of significant difference between negative controls and experimental group (Student’s t-test: *P<0.05). **(g)** and **(h)** Schematic diagram of constructs for effectors and reporters (upper panel), and mean (±SE) relative NaACS1_pro_::LUC **(g)** and NaACO1_pro_::LUC **(h)** transient transcription activity activated by NaWRKY3 in *N. benthamiana* plants (lower panel). The *Agrobacterium* with NaACS1_pro_::LUC **(g)** and NaACO1_pro_::LUC **(h)** reporter was co-injected with those with indicated effector into leaves of *N. benthamiana*. After 36 h, values represented the NaACS1_pro_::LUC **(g)** and NaACO1_pro_::LUC **(h)** activity relative to the internal control (35S::REN activity) were obtained. Asterisks indicated significant differences between control and Ov-NaWRKY3 samples (Student’s *t*-test; n=4; *P<0.05, ***P<0.005).

Indeed, several ethylene synthetic enzyme genes were found to be decreased in infected irWRKY3 plants during transcriptome analysis. qPCR results showed *A. alternata*-induced *NaACS1* and *NaACO1* were strongly impaired in irWRKY3 plants (Fig. 4a). EMSA results indicated that NaWRKY3 protein could specifically bind to the probes designed from the W-boxes of *NaACS1* and *NaACO1* promoters (Fig. 4b-d). Further ChIP-qPCR assay showed that the DNA from *NaACS1* and *NaACO1* regions around the EMSA probes were more enriched by anti-GFP, indicating NaWRKY3 can bind directly to the promoters of *NaACS1* and *NaACO1 in vivo* (Fig. 4e and 4f).

To determine whether NaWRKY3 was able to activate *NaACS1* and *NaACO1* promoter, a dual-LUC reporter assay was employed. As expected, the expression of the LUC reporter driven by NaACS1_pro_ and NaACO1_pro_ were activated by NaWRKY3 in leaves of *N. benthamiana* plants (Fig. 4g and 4h).

Thus, our data demonstrates that NaWRKY3 acts as a positive regulator of *A. alternata*-induced ethylene by directly binding to *NaACS1* and *NaACO1* promoter and activating their expression.

### NaWRKY3 is required for *A. alternata*-induced phytoalexins

Scopoletin and scopolin are phytoalexins in *N. attenuata* regulated by JA and ethylene signaling pathways for *A. alternata* resistance (Sun et al., 2014; Sun et al., 2017). The levels of scopoletin and scopolin accumulated in irWRKY3 plants were greatly reduced at both 1 and 3 dpi (Fig. 5a). Similarly, *A. alternata*-induced expression of *NaF6’H1*, the key enzyme gene of the scopoletin biosynthesis, was also strongly impaired in the irWRKY3 plants (Fig. 5a).

**Fig. 5.**
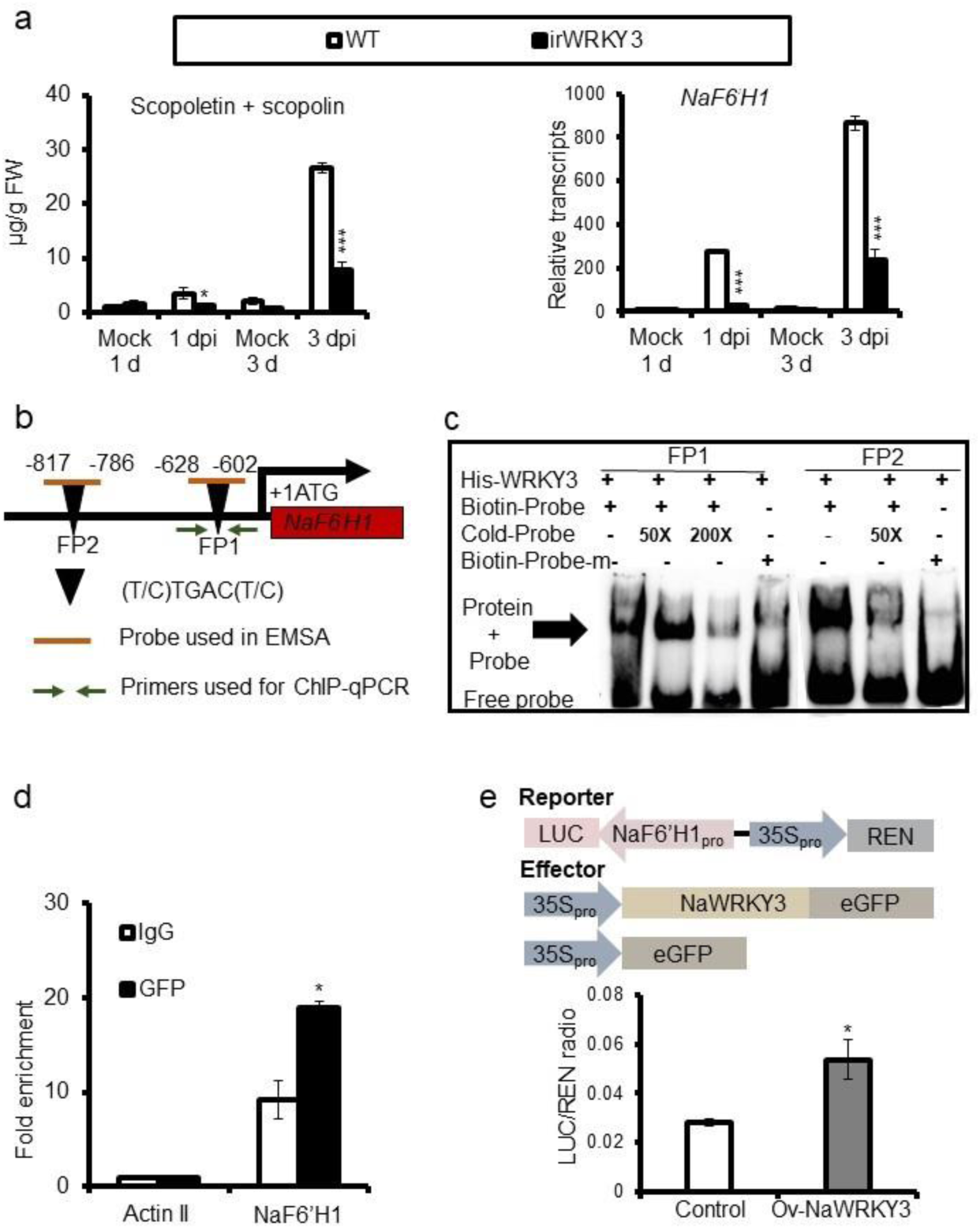
NaWRKY3 is required for scopoletin and scopolin accumulation and the expression of *NaF6’H1* after *A. alternata* inoculation. **(a)** Mean (±SE) relative *A. alternata*-induced scopoletin and scopolin levels and *NaF6’H1* transcripts in five replicate 0 leaves of WT and irWRKY3 plants. Asterisks indicated the levels of significant differences between WT and irWRKY3 plants with the same treatments (Student’s *t*-test: *P<0.05, ***P<0.005). **(b)** Schematic diagram of the promoter of *NaF6’H1*. Black triangles indicated the W-box motifs. Short orange lines indicated DNA probes used for EMSA, and green arrows indicated primers used for ChIP-qPCR assays. The translational start site (ATG) was shown at position +1. **(c)** EMSA showed that His-NaWRKY3 could bind to the promoter of *NaF6’H1* directly. 50- and 200-fold excess of unlabeled probes were used for competition. Biotin-Probe-m was the probe with mutated W-box motif. **(d)** Mean (±SE) fold enrichment on the NaWRKY3-eGFP plants in ChIP-qPCR. DNA from NaWRKY3-eGFP over-expressed plants at 1 dpi was immune-precipitated by GFP or IgG antibody, and was further analyzed by real time PCR with indicated primers. The immune-precipitated DNA from IgG antibody was served as control, and asterisks indicated levels of significant difference between negative controls and experimental group (Student’s t-test: *P<0.05). **(e)** Schematic diagram of constructs for effectors and reporters (upper panel), and mean (±SE) relative NaF6’H1_pro_::LUC transient transcription activity activated by NaWRKY3 in *N. benthamiana* plants (lower panel). The *Agrobacterium* with NaF6’H1_pro_::LUC reporter was co-injected with those with indicated effector into leaves of *N. benthamiana*. After 36 h, values represented the NaF6’H1pro::LUC activity relative to the internal control (35S::REN activity) were obtained. Asterisks indicated significant differences between control and Ov-NaWRKY3 samples (Student’s *t*-test; n=4; *P<0.05).

EMSA results indicated that NaWRKY3 could specifically bind to two probes designed from the W-boxes in the promoter region of *NaF6’H1* (Fig. 5b and 5c). This binding of NaWRKY3 to the promoter of *NaF6’H1 in vivo* was further verified by ChIP-qPCR assay (Fig. 5d). Dual-LUC assay results showed that NaWRKY3 could significantly induced the LUC reporter expression which was driven by the promoter of *NaF6’H1* (Fig. 5e).

Our data demonstrates that NaWRKY3 is required for *A. alternata*-induced scopoletin accumulation by directly binding to *NaF6’H1* promoter and activating its expression.

### Regulation of *NaRboh D*-dependent defensive responses by NaWRKY3

In *N. attenuata*, NaRboh D had been reported to be involved in the ROS burst after insect herbivore attack (Wu et al., 2013). Silencing *NaRboh D* (Fig. 6a) also strongly reduced ROS production after *A. alternata* inoculation at 12 hpi (Supplemental Fig. 6). Interestingly, *A. alternata* infection usually leads to stomata closure, which is essential for plant resistance to this fungus (Sun et al., 2014). However, *A. alternata*-induced stomata closure was abolished in *NaRboh D*-silenced irRboh D plants (Fig. 6a). Consistently, irRboh D plants were highly susceptible to *A. alternata* (Fig. 6b).

**Fig. 6.**
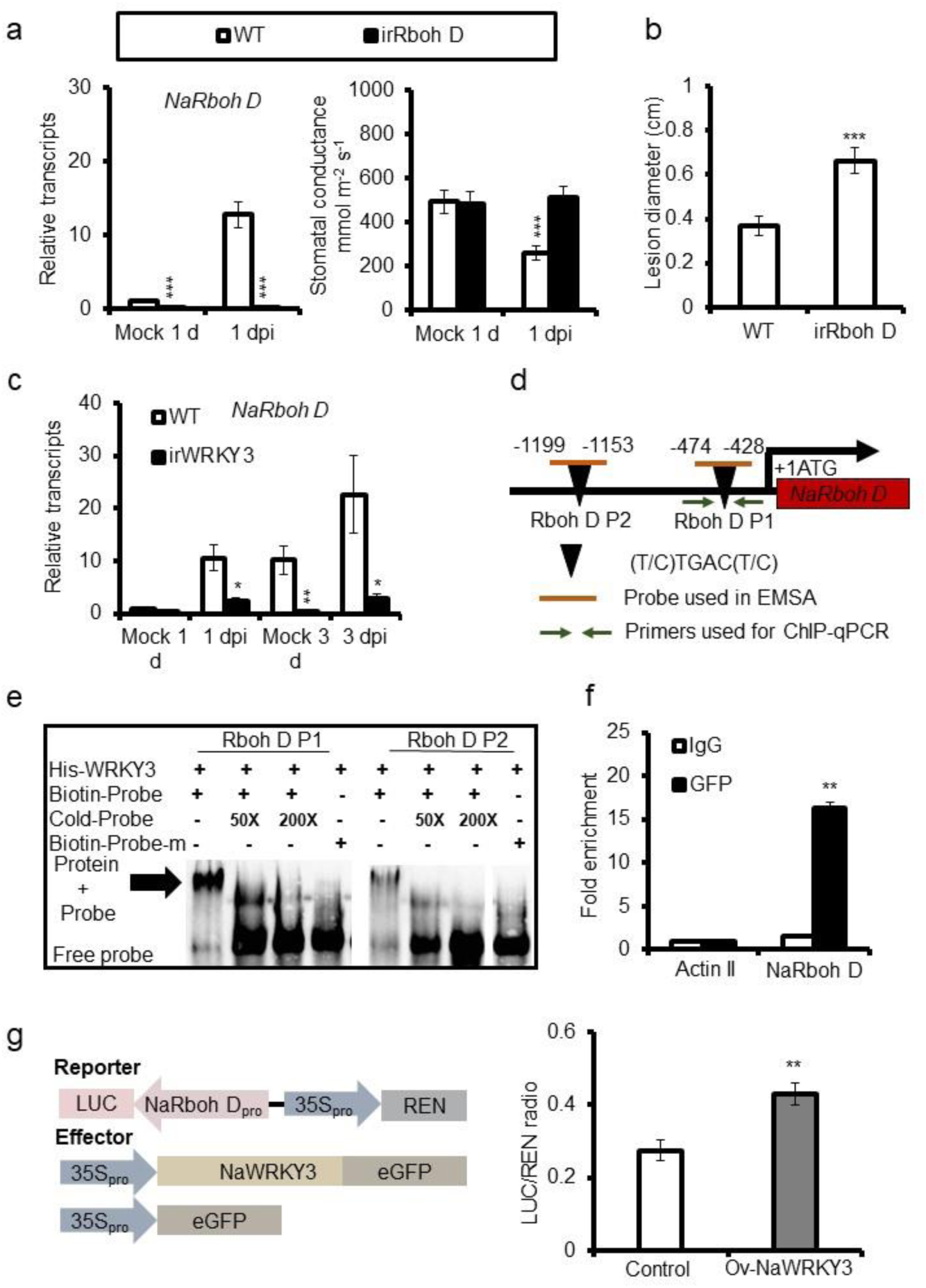
NaWRKY3 is required for the expression of *A. alternata* resistance *NaRboh D*. **(a)** Mean (±SE) relative *A. alternata*-induced *NaRboh D* transcripts (left panel) and stomatal conductance (right panel) as measured in five replicate 0 leaves at 1 dpi. Asterisks indicated levels of significant difference between WT and irRboh D samples (Student’s *t*-test: ***P<0.005). **(b)** Mean (±SE) diameter of necrotic lesions of 20 inoculation sites on 0 leaves from WT and irRboh D plants. The lesion area analyzes at 4 dpi. Asterisks indicated significant difference from the WT and irRboh D plants according to Student’s *t*-test at ***P < 0.005. **(c)** Mean (±SE) relative *A. alternata*-induced *NaRboh D* transcripts as measured by qPCR in five replicate leaves of WT and irWRKY3 plants at 1 and 3 dpi. Asterisks indicated the levels of significant differences between WT and irWRKY3 plants with the same treatments (Student’s *t*-test: *P<0.05, **P<0.01). **(d)** Schematic diagram of the promoter of *NaRboh D*. Black triangles indicated the W-box motifs. Short orange lines indicated DNA probes used for EMSA, and green arrows indicated primers used for ChIP-qPCR assays. The translational start site (ATG) was shown at position +1. **(e)** EMSA showed that His-NaWRKY3 could bind to the promoter of *NaRboh D* directly. 50- and 200-fold excess of unlabeled probes were used for competition. Biotin-Probe-m was the probe with mutated W-box motif. **(f)** Mean (±SE) fold enrichment on the NaWRKY3-eGFP plants in ChIP-qPCR. DNA from NaWRKY3-eGFP over-expressed plants at 1 dpi was immune-precipitated by GFP or IgG antibody, and was further analyzed by real time PCR with indicated primers. The immune-precipitated DNA from IgG antibody was served as control, and asterisks indicated levels of significant difference between negative controls and experimental group (Student’s t-test: **P<0.01). **(g)** Schematic diagram of constructs for effectors and reporters (left panel), and mean (±SE) relative NaRboh D_pro_::LUC transient transcription activity activated by NaWRKY3 in *N. benthamiana* plants (right panel). The *Agrobacterium* with NaRboh D_pro_::LUC reporter was co-injected with those with indicated d effector into leaves of *N. benthamiana*. After 36 h, values represented the NaRboh Dpro::LUC activity relative to the internal control (35S::REN activity) were obtained. Asterisks indicated significant differences between control and Ov-NaWRKY3 samples (Student’s *t*-test; n=4; **P<0.01).

We found that *A. alternata*-induced *NaRboh D* transcripts were strongly decreased in irWRKY3 plants during transcriptome analysis. qPCR results confirmed that silencing *NaRboh D* greatly impaired *A. alternata*-induced *NaRboh D* expression (Fig. 6c). EMSA and ChIP-qPCR results showed that NaWRKY3 could specifically bind to the promoter of *NaRboh D in vitro* and *in vivo* (Fig. 6d-f). Dual-LUC experiments further showed that NaWRKY3 activated LUC expression driven by *NaRboh D* promoter (Fig. 6g).

Thus, our data suggests that NaWRKY3 positively regulates *NaRboh D* expression through binding to its promoter to activate its expression. Meanwhile, *NaRboh D* is involved in the resistance to *A. alternata* via ROS and stomata closure.

### NaWRKY3 is required for the expression of *NaBBL28* after *A. alternata* inoculation

Berberine bridge like (BBL) genes are believed to encode enzymes involved in specialized metabolism such as alkaloids biosynthesis in *Nicotiana*. There were several BBL genes found to be induced by *A. alternata* in WT plants (Supplemental Fig. 4a and 4b, Fig. 7a). Among them, *NaBBL28* (XM_019378340.1) transcripts were strongly increased in response to *A. alternata*, while in irWRKY3 plants the *NaBBL28* transcripts were dramatically decreased (Fig. 7a). Furthermore, plants with silenced *NaBBL28* via VIGS were highly susceptible to *A. alternata* (Supplemental data 4c and 4d).

**Fig. 7.**
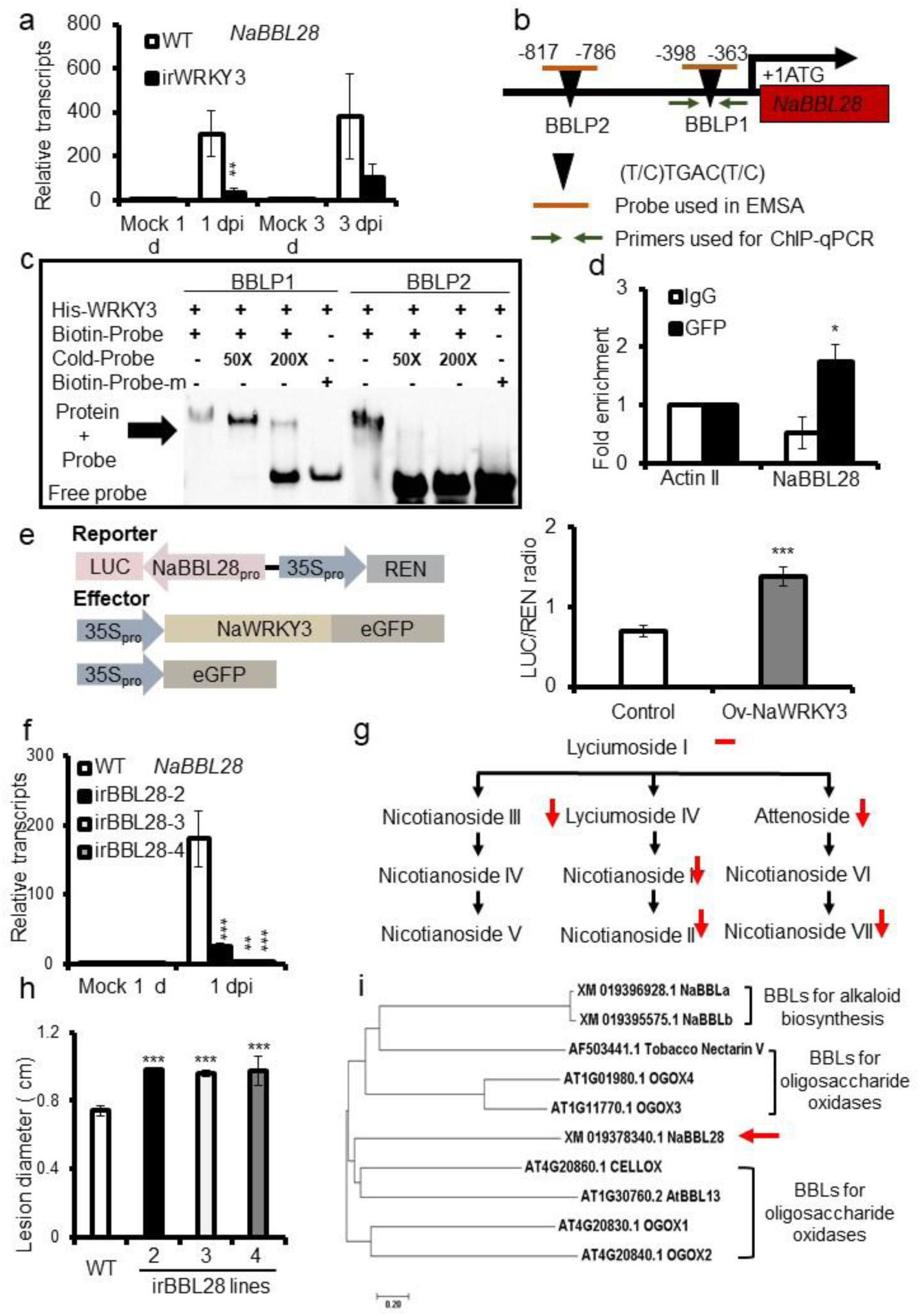
NaWRKY3 is required for the expression of *A. alternata* resistance *NaBBL28*. **(a)** Mean (±SE) relative *A. alternata*-induced *NaBBL28* transcripts as measured by qPCR in five replicate leaves of WT and irWRKY3 plants at 1 and 3 dpi. Asterisks indicated the levels of significant differences between WT and irWRKY3 plants with the same treatments (Student’s *t*-test: **P<0.01). **(b)** Schematic diagram of the promoter of *NaBBL28*. Black triangles indicated the W-box motifs. Short orange lines indicated DNA probes used for EMSA, and green arrows indicated primers used for ChIP-qPCR assays. The translational start site (ATG) was shown at position +1. **(c)** EMSA showed that His-NaWRKY3 could bind to the promoter of *NaBBL28* directly. 50- and 200-fold excess of unlabeled probes were used for competition. Biotin-Probe-m was the probe with mutated W-box motif. **(d)** Mean (±SE) fold enrichment on the NaWRKY3-eGFP plants in ChIP-qPCR. DNA from *NaWRKY3-eGFP* over-expressed plants at 1 dpi was immune-precipitated by GFP or IgG antibody, and was further analyzed by real time PCR with indicated primers. The immune-precipitated DNA from IgG antibody was served as control. Asterisks indicated the levels of significant differences between WT and irWRKY3 plants with the same treatments (Student’s t-test: *P<0.05). **(e)** Schematic diagram of constructs for effectors and reporters (left panel), and mean (±SE) relative NaBBL28_pro_::LUC transient transcription activity activated by NaWRKY3 in *N. benthamiana* plants (right panel). The *Agrobacterium* with NaBBL28_pro_::LUC reporter was co-injected with those with indicated effector into leaves of *N. benthamiana*. After 36 h, values represented the NaBBL28pro::LUC activity relative to the internal control (35S::REN activity) were obtained. Asterisks indicated significant differences between control and Ov-NaWRKY3 samples (Student’s *t*-test; n=4; ***P<0.005). **(f)** Mean (±SE) relative *A. alternata*-induced *NaBBL28* transcripts as measured by qPCR in five replicate 0 leaves of WT and irBBL28 (irBBL28-2, irBBL28-3, irBBL28-4) plants at 1 dpi. Asterisks indicated the levels of significant differences between WT and irBBL28 plants with the same treatments (Student’s *t*-test: **P<0.01, ***P<0.005). **(g)** HGL-DTGs biosynthetic pathway was used for indication of reduced HGL-DTGs. Red downward pointing arrows indicated the levels of HGL-DTGs significant decreased in irBBL28 leaves, short red lines indicated the levels of Lyciumoside I were not altered between the WT and irBBL28 plants. **(h)** Mean (±SE) diameter of necrotic lesions of 20 inoculation sites on 0 leaves from WT and irBBL28 plants. The lesion area analyzes at 4 dpi. Asterisks indicated significant difference from the WT and irBBL28 plants according to Student’s *t*-test at ***P < 0.005. (i) Phylogenetic analysis of NaBBL28 and other BBL proteins. BBLs which have been functionally characterized as oligosaccharide oxidase and alkaloid biosynthesis enzymes in *Arabidopsis* and *Nicotiana* were used for alignment. NaBBL28 was indicated with the red arrow.

EMSA showed that NaWRKY3 could bind to two W-boxes in the promoter region of *NaBBL28 in vitro*, and this binding was further verified by ChIP-qPCR assay (Fig. 7b-d). Dual-LUC assay was employed to show that NaWRKY3 could significantly induce the LUC activity driven by the promoters of *NaBBL28* (Fig. 7e).

To further investigate the role of *NaBBL28* in *N. attenuata* resistance to *A. alternata*, stable *NaBBL28* RNAi plants were obtained (Fig. 7f). qPCR showed *NaBBL28* expression was successfully silenced in three *NaBBL28* RNAi lines, irBBL28-2, irBBL28-3 and irBBL28-4 after *A. alternata* inoculation at 1 dpi (Fig. 7f). All three RNAi lines were highly susceptible to *A. alternata* (Fig. 7h), which was consistent with the phenotype observed in VIGS plants (Supplemental Fig. 4c and 4d).

NaBBL28 had a FAD (flavin adenine dinucleotide)-binding domain that was known feature of BBE-like proteins, belonged to FAD/FMN-containing dehydrogenase family with a N-terminal signal peptide (Supplemental Fig. 4e and 4f). A phylogenetic analysis including functional characterized BBLs (Carter and Thornburg, 2004; Benedetti et al., 2018; Locci et al., 2019) showed a closer relationship of NaBBL28 to oxidize glucose/oxidize cellulose rather than alkaloid-related BBLs (Fig. 7g), suggesting that *NaBBL28* might act as a oligosaccharide oxidase or dehydrogenase.

*N. attenuata* plants accumulated large quantities of defensive 17-hydroxygeranyllinalool diterpene glycosides (HGL-DTGs). We thus investigated HGL-DTGs in *NaBBL28* RNAi lines by HPLC-MS/MS. Silencing *NaBBL28* resulted several HGL-DTGs decreased significantly. A marked reduction in Nicotianoside III, Attenoside, Nicotianosides I, Nicotianoside II, Nicotianoside VII were observed in *NaBBL28*-silenced plants, whereas little effect was found in their precursor Lyciumoside I (Supplemental Fig. 5). The results revealed that NaBBL28 might be important to hydroxylate the aglycones especially the rhamnose at C-3 of Lyciumoside I to form Nicotianoside III, and Attenoside.

Taken together, our results reveal that NaWRKY3 positively regulates *NaBBL28*, which is required for *A. alternata* resistance likely by acting as a rhamnose hydroxylase of HGL-DTGs.

### NaWRKY3 negative regulates its own expression after *A. alternata* inoculation

To further characterize the role of NaWRKY3 in *A. alternata* resistance, we generated transgenic *N. attenuata* plants with constitutively over-expressed *NaWRKY3* in two versions (Supplemental Fig. 1c). Under the control of 35S promoter, *NaWRKY3* ORF with 5’-UTR (OV3C1) and genomic DNA of NaWRKY3 (OV3G1) were fused with eGFP and transformed into *N. attenuata* plants. qPCR and Western Blot showed that the *NaWRKY3-eGFP* transcripts and proteins were successfully expressed in both versions of transgenic plants (Supplemental Fig. 1b and 1c). However, we could not detect the higher levels of total *NaWRKY3* transcripts in all over-expressed lines (Supplemental Fig. 1c).

Both EMSA and ChIP-qPCR indicated that NaWRKY3 could specifically bind to its own promoter (Fig. 8a-c). Furthermore, NaWRKY3 could significantly repress the LUC activity driven by its own promoter in dual-LUC assay, and this suppression could not be relieved even at *A. alternata* infected condition (Fig. 8d).

**Fig. 8.**
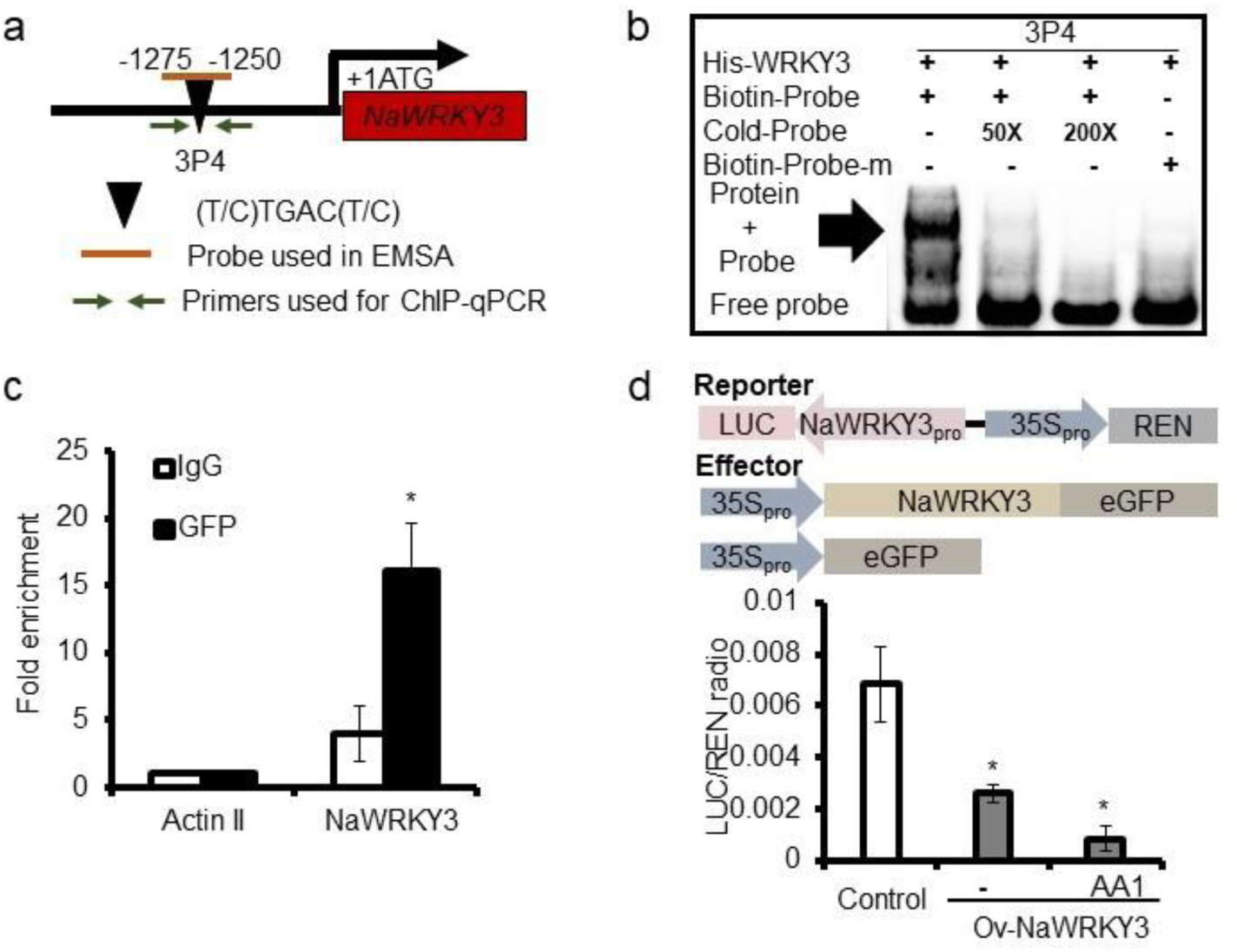
NaWRKY3 negative regulates the expression of itself after *A. alternata* inoculation. **(a)** Schematic diagram of the promoter of *NaWRKY3*. Black triangle indicated the W-box motif. Short orange line indicated DNA probes used for EMSA, and green arrows indicated primers used for ChIP-qPCR assays. The translational start site (ATG) was shown at position +1. **(b)** EMSA showed that His-NaWRKY3 could bind to the promoter of *NaWRKY3* directly. 50- and 200-fold excess of unlabeled probes were used for competition. Biotin-Probe-m was the probe with mutated W-box motif. **(c)** Mean (±SE) fold enrichment on the NaWRKY3-eGFP plants in ChIP-qPCR. DNA from NaWRKY3-eGFP over-expressed plants at 1 dpi was immune-precipitated by GFP or IgG antibody, and was further analyzed by real time PCR with indicated primers. The immune-precipitated DNA from IgG antibody was served as control. Asterisks indicated significant differences between control and Ov-NaWRKY3 samples (Student’s t-test; n=4; *P<0.05). **(d)** Schematic diagram of constructs for effectors and reporters (left panel), and mean (±SE) relative NaWRKY3_pro_::LUC transient transcription activity activated by NaWRKY3 in *N. benthamiana* plants (right panel). The *Agrobacterium* with NaWRKY3_pro_::LUC reporter was co-injected with those with indicated effector into leaves of *N. benthamiana*. AA1 indicated *A. alternata* inoculation 1 dpi, asterisks indicated significant differences between control and Ov-NaWRKY3 samples (Student’s *t*-test; n=4; *P<0.05).

Thus, we find that NaWRKY3 could bind to its own promoter but acts as a transcriptional repressor.

## Discussion

### NaWRKY3 regulates phytoalexins and upstream signals

Many WRKYs have been reported to be involved in plant resistance to pathogens, including AtWRKY33, AtWRKY8, AtWRKY57, AtWRKY46, AtWRKY70, and AtWRKY53 in *Arabidopsis* (Birkenbihl et al., 2012; Chen et al., 2018; Jiang and Yu, 2016; Hu et al., 2012). AtWRKY46 coordinates with AtWRKY70 and AtWRKY53 in basal resistance against *P. syringae* partially involved in SA-signaling pathway (Hu et al., 2012). AtWRKY8 is a negative regulator of basal resistance to *P. syringae* due to decrease expression of *PR1*and positive regulator to *B. cinerea* by elevating *PDF1.2* transcripts (Chen et al., 2010; Chen et al., 2018). NbWRKY7, 8, 9 and 11 played redundant roles in immunity to *P. infestans* (Adachi et al., 2015). *A. alternata* is notorious necrotrophic fungal pathogen which causes severe loss in *Nicotiana* species. However, to date no WRKYs has been identified to be involved in defense responses to this pathogen.

Previously, we have shown that *NaF6’H1* is the key enzyme gene for the biosynthesis of phytoalexins scopoletin and scopolin, which serves as the first line of defense to *A. alternata* in *N. attenuata* (Sun et al., 2014). Analysis of *NaF6’H1* promoter revealed that its expression was likely regulated by some WRKYs as several W-boxes occurred in the promoter region. Indeed, several WRKYs were found to be highly induced by *A. alternata* during transcriptome analysis (Song et al., 2019). Among them, *NaWRKY3* was one of the WRKYs elicited significantly by *A. alternata* not only during transcriptome analysis but also verified by real time PCR (Fig. 1). Importantly, silencing NaWRKY3 resulted plants highly susceptible to *A. alternata* (Fig. 1). Further investigation revealed that NaWRKY3 was required for *A. alternata*-induced scopoletin and scopolin accumulation by directly binding to *NaF6’H1* promoter and activating its expression, which were proved by EMSA, ChIP-qPCR, dual-LUC assay and analysis of RNAi plants (Fig. 5). Thus, NaWRKY3 was the first WRKY identified in *Nicotiana* species to regulate plant resistance to *A. alternata* by regulation of biosynthesis of phytoalexins, scopoletin and scopolin.

Plant hormones JA and ethylene are usually associated with plant resistance to necrotrophs (Glazebrook, 2005). *N. attenuata* plants activate both JA and ethylene signaling after *A. alternata* attack, which subsequently regulate scopoletin and scopolin biosynthesis to defend against this fungus (Sun et al., 2014; Sun et al., 2017). Here we reported that NaWRKY3 not only regulated phytoalexin biosynthesis directly through *NaF6’H1* (Fig. 5), but also controlled *A. alternata*-induced levels of JA and ethylene through binding and activating *NaLOX3*, *NaACS1* and *NaACO1* promoter (Fig. 2, Fig. 4).

Rboh D is an NADPH oxidase, which is crucial for ROS production after pathogens infection, and is essential for plant resistance to pathogens, AtWRKY55 activates the expression of *AtRboh D* positively regulates defense against *P. syringae* (Wang et al., 2020). In this study, we found that *NaRboh D* was involved in the resistance to *A. alternata* via ROS and stomata closure (Fig. 6 and Supplemental Fig. 6). Interestingly, NaWRKY3 positively regulated *NaRboh D* expression through binding to its promoter to activate its expression (Fig. 6).

Thus, we propose that NaWRKY3 is a regulator of defense networks in *N. attenuata* to *A. alternata*; it not only regulates the direct defense chemicals scopoletin and scopolin, but also controls upstream signals including JA, ethylene and NaRboh D-based ROS.

### NaWRKY3 is a functional homolog of AtWRKY33, but with different working mechanism

In *Arabidopsis*, *AtWRKY33* is proposed as a key regulator of hormonal and metabolic responses in plant resistance to *B. cinerea* (Birkenbihl et al., 2012; Zhou et al., 2022). Our NaWRKY3 shared 46.44% amino acid sequence identity to AtWRKY33, and was the most closed AtWRKY33 homolog in *N. attenuata* (Supplemental Fig. 7).

To some extent, AtWRKY33 and NaWRKY3 act in a similar way in defense responses but with different target genes. *AtWRKY33* was involved in *B. cinerea*-induced ethylene produced by regulation of *AtACS2* and *AtACS6* expression through directly binding to the promoter of those genes (Li et al., 2012). Similarly, NaWRKY3 also acted as a positive regulator of *A. alternata*-induced ethylene, it could directly bind to *NaACS1* and *NaACO1* promoter and activate their expression (Fig. 4). AtWRKY33 bound to *PAD3* promoter and positively regulated phytoalexin camalexin biosynthesis (Zhou et al., 2022). Scopoletin and scopolin were phytoalexins produced in *N. attenuata* which were required for *A. alternata* resistance (Sun et al., 2014; Sun et al., 2017). *A. alternata*-induced *NaF6’H1* expression was strongly reduced in *NaWRKY3*-silenced plants, and *NaF6’H1* was regulated by NaWRKY3 by directly binding to its promoter and activating its expression (Fig. 5).

More interestingly, some of the target genes or pathways are regulated by AtWRKY33 and NaWRKY3 in opposite ways. *AtRboh D* expression was elevated in *Arabidopsis* upon *B. cinerea* infection, but its levels in *wrky33* mutants were similar to those of WT plants (Birkenbihl et al., 2012). However, *A. alternata*-induced *NaRboh D* expression was greatly impaired in *NaWRKY3*-silenced plants and we demonstrated that NaWRKY3 positively regulated *NaRboh D* through binding to its promoter to activate its expression after inoculation of *A. alternata* (Fig. 6). At 1 dpi, *wrky33* mutants accumulated more *B. cinerea*-induced JA levels than WT, which might be due to AtWRKY33 negatively regulated SA signaling in *Arabidopsis* (Birkenbihl et al., 2012). But in *N. attenauta*, *A. alternata*-induced JA levels were significantly reduced in *NaWRKY3*-silenced plants, and we found that JA biosynthetic gene *NaLOX3* was the direct target of NaWRKY3 (Fig. 2).

Thus, we propose that NaWRKY3 is a functional homolog of AtWRKY33. Although NaWRKY3 and AtWRKY33 are both required for plant resistance to necrotrophic fungal pathogens and involved in phytohormone signaling and phytoalexin biosynthesis, their detailed mode of action is quite different.

### NaWRKY3 regulates *NaBBL28*, a gene required for *N. attenuata* resistance to *A. alternata* and related to HGL-DTGs oligosaccharide hydroxylation

The *BBL* genes are believed to catalyze the oxidative cyclization in specialized metabolism biosynthesis. Overexpressing of *BBL* genes *OGOX1* and *CELLOX* in *A. thaliana* increased plant resistance to *B. cinerea*, because oxidized oligogalacturonides (OGs) and cellodextrins (CDs) are inefficient carbon source that cannot be utilized by *B. cinerea* (Benedetti et al., 2018; Locci et al., 2019). However, whether any *BBL* genes are involved in defense responses to *A. alternata* is unknown. In our study, we found that *NaBBL28* gene was strongly up-regulated in response to *A. alternata* inoculation at both 1 and 3 dpi in 0 leaves (Fig. 7), and bigger lesions were observed in stable *NaBBL28* RNAi plants and *NaBBL28* VIGS plants (Fig. 7h, Supplemental Fig. 4d). These findings suggest that *NaBBL28* is required for the resistance of *N. attenuata* to *A. alternata*.

Currently, there is not report how BBLs are regulated. In our study, we found several W-boxes in the promoter of *NaBBL28* and strongly decreased transcripts of *NaBBL28* in irWRKY3 plants (Fig. 7). Further EMSA, ChIP-qPCR and dual-LUC assay demonstrated NaWRKY3 directly up-regulated the expression of *NaBBL28* by binding and activating its promoter (Fig. 7b-e).

The substrates of NaBBL28 are still unclear. BBLs acting as glycosides oxidases is proved in PpBBE1, AtBBE13, Nectarin V, OGOX1, OGOX2, OGOX3, OGOX4, CELLOX (Carter and Thornburg, 2004; Benedetti et al., 2018; Toplak et al., 2018; Locci et al., 2019; Scortica et al., 2022). HGL-DTGs are acyclic diterpene glycosides found in *Nicotiana* species (Heiling et al., 2016). HGL-DTGs have a 17-hydroxygeranyllinalool (HGL) backbone that can be decorated at the C-3 and C-17 positions with glucose and rhamnose by oligosaccharide oxidases and dehydrogenases (Heiling et al., 2016), forms the precursor Lyciumoside I. Sequence alignment indicated that NaBBL28 was more likely an oligosaccharide oxidase or dehydrogenase (Fig. 7i, Supplemental Fig. 4e and 4f). In our study, the levels of precursor Lyciumoside I did not alter in *NaBBL28*-silenced plants, while Nicotianoside III, Attenoside, Nicotianosides I, Nicotianoside II, and Nicotianoside VII were all significantly decreased in *NaBBL28*-silenced plants (Fig. 7 and Supplemental Fig. 5), suggesting that NaBBL28 might be a rhamnose dehydrogenase, adding rhamnose to C-3 of Lyciumoside I.

It’s for the first time that a WRKY was identified in the transcriptional regulation of *BBL* gene. NaWRKY3 positively regulates *NaBBL28*, which is required for *A. alternata* resistance. NaBBL28 acts likely as an oligosaccharide dehydrogenase participating in HGL-DTGs hydroxylation at C-3 of the Lyciumoside I aglycones. However, more work is still needed to prove whether Nicotianoside III, Attenoside, Nicotianosides I, Nicotianoside II, and Nicotianoside VII are phytoalexins against *A. alternata* or not.

### The lncRNA *L2* required for *A. alternata* resistance is regulated by NaWRKY3 and influences JA accumulation

Plant lncRNAs have been reported to play important roles pathogen resistance. *GblncRNA7* and *GblncRNA2* regulate the resistance to *V. dahlia* in opposite ways (Zhang et al., 2022). LncRNA33732, Sl-lncRNA15492 and Sl-lncRNA39896 are important regulators for tomato resistance to *P. infestans* (Cui et al., 2019; Jiang et al., 2020; Hong et al., 2022). However, evidence regarding any lncRNAs involved in defense responses to *A. alternata* is still lacking. The presented data here strongly indicated that *N. attenuata* plants up-regulated lncRNA *L2* as a defense regulator against *A. alternata*, since its levels were dramatically increased after *A. alternata* infection (Fig. 3c), and bigger lesions were developed in *L2*-silenced plants via RNAi or VIGS (Fig. 3 and Supplemental Fig. 3). Further analysis revealed that silencing *L2* impaired *A. alternata*-induced JA and *NaF6’H1* expression (Fig. 3). However, the detail mechanism how lncRNA *L2* regulates these responses is currently needed more investigation.

### NaWRKY3 is a master regulator of defense networks in *N. attenuata* to *A. alternata*

In this study, we demonstrate that NaWRKY3 is a master regulator of defense network against *A. alternata* in *N. attenuata*. Here is the working model we proposed (Fig. 9). When host plants sense the attack of *A. alternata*, NaWRKY3 up-regulates 1) JA biosynthesis by regulation of *NaLOX3* and lncRNA *L2*; 2) ethylene biosynthesis by *NaACS1* and *NaACO1*; and 3) the respiratory burst oxidase *NaRboh D*, which is required for *A. alternata*-induced ROS and stomal closure. These results suggest that NaWRKY3 regulates up-stream signaling including JA, ethylene and ROS. In addition, NaWRKY3 also regulates downstream specialized metabolites involved in *A. alternata* resistance. Scopoletin and scopolin biosynthesis is directly activated by NaWRKY3 on *NaF6’H1* expression, and NaWRKY3 also controls NaBBL28, which is required for *A. alternata* resistance and likely involved in hydroxylation of the rhamnose of Lyciumoside I. Finally, as a key transcriptional regulator, the expression of *NaWRKY3* is fine-tuned by itself during defense.

**Fig. 9.**
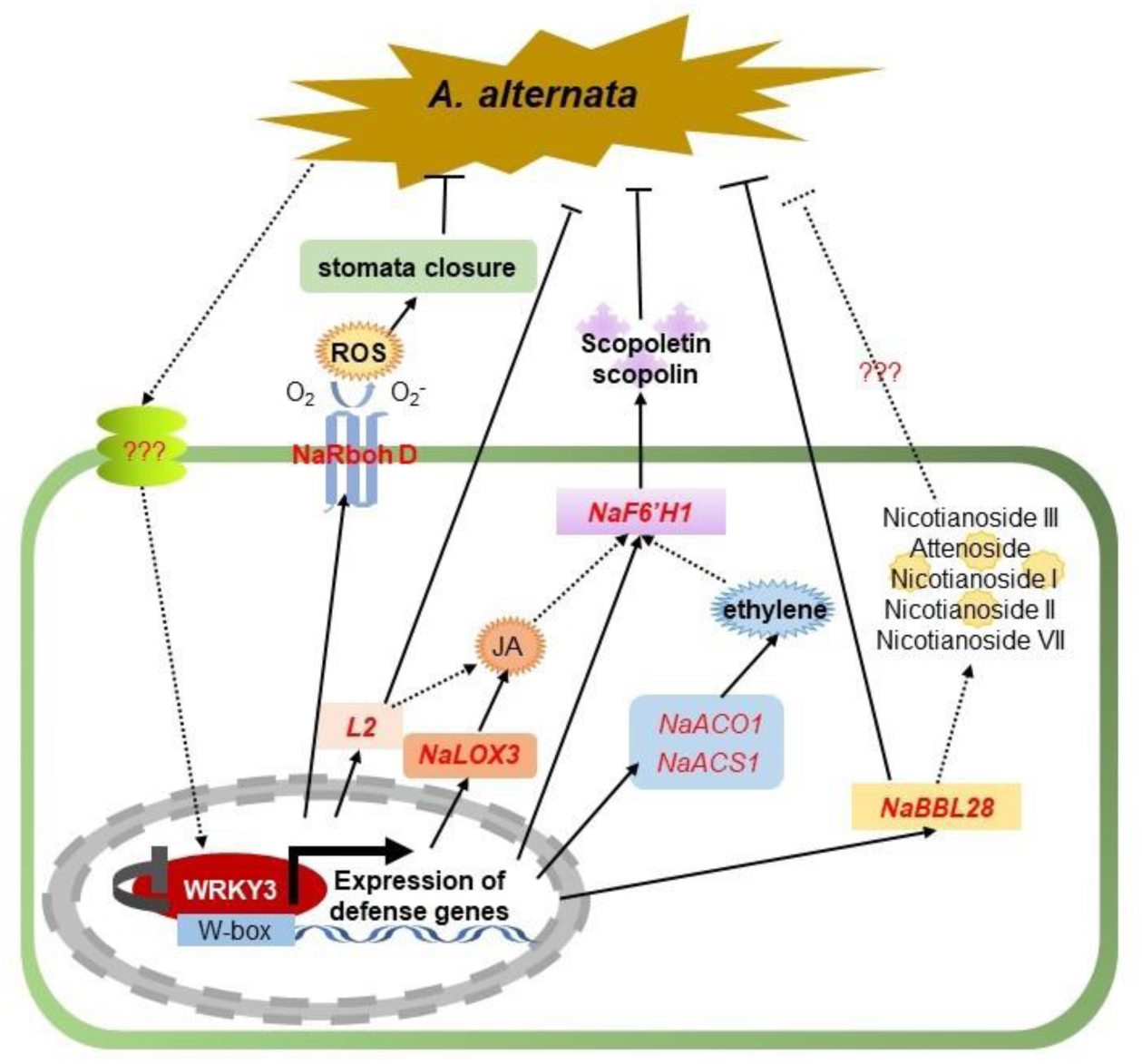
Working model of the defense network regulated byNaWRKY3 in *N. attenuata* after *A. alternata* challenge. *N. attenuata* plants increase *NaWRKY3* expression after *A. alternata* infection. The NaWRKY3 protein then binds to the promoters of jasmonic acid (JA) synthetic gene *NaLOX3*, JA related LncRNA *L2*, ethylene synthetic gene *NaACS1* and *NaACO1*, respiratory burst oxidase gene *NaRboh D*, scopoletin and scopolin synthetic gene *NaF6’H1* and HGL-DTGs related dehydrogenase *NaBBL28*, and activates their transcription, thereby regulating a complicate defense network against *A. alternata*. It includes defense hormones JA and ethylene, NaRboh D-based ROS for stomata closure, scopoletin and scopolin as the direct chemical weapon, LncRNA *L2* for JA and *NaF6’H1* expression, and NaBBL28 for hydroxylation of the rhamnose of Lyciumoside I to form Nicotianoside III, Attenoside, and other HGL-DTGs. Silencing LncRNA *L2*, *NaBBL28* and *NaRboh D* separately will lead to plants susceptible to the fungus. Finally, NaWRKY3 fine-tuned its expression by repressing itself.

## Materials and Methods

### Plant growth and generation of stably transformed plants

Seeds of 33^rd^ inbreeding lines of *N. attenuata* were used as the wild-type (WT) in all experiments. Stably transformed lines of irWRKY3 (*NaWRKY3*-deficient; Skibbe et al., 2008), irRboh D (*NaRboh D*-deficient; Wu et al., 2013) were kindly provided by Prof. Ian T Baldwin.

Stable transgenic plants of irBBL28 (*NaBBL28*-RNAi), irL2 (*L2*-RNAi), OV3G1 (over-expression of *NaWRKY3* ORF with 5’-UTR), OV3C1 (over-expression of genomic *NaWRKY3*) were generated in this study as followed. Seed germination and plant growth were conducted as described by Krügel et al. (2002).

### Generation of stably transformed plants

For RNAi lines, inverted-repeat orientation of the target gene fragments of lncRNA *L2* or *NaBBL28* were amplified by primers (Supplementary table) and inserted into the pRESC8. For over-expression line, *NaWRKY3* ORF with 5’-UTR (OV3C1) and genomic *NaWRKY3* (OV3G1) was infused with eGFP and inserted into pCAMBIA1301 under the control of 35S promoter. These constructed vectors were transformed into *N. attenuata* WT plants by using *Agrobacterium*-mediated transformation procedure (Zhao et al., 2021). Single-insertion RNAi lines of irL2 (irL2-1, irL2-3) and irBBL28 (irBBL28-2, irBBL28-3, irBBL28-4) were identified, bred to homozygosity in T2 generation and used in this study.

### Total RNA extraction, RT-qPCR analysis and RNA-seq

Total RNA was extracted from a 1.5 ×1.5 cm2 area of leaf lamina with the inoculation site at the center using TRIzol reagent. Around 1 μg of total RNA was used as templates for reverse transcription into cDNA with reverse transcriptase (www.thermofisher.cn). qPCR was performed using SYBR Green mix (BioRad) on a CFX Connect qPCR System (Bio-Rad) and gene-specific primers according to the manufacturer’s instructions. For RT-qPCR analysis, the cDNA was diluted 1.5 times, and 2 μL reaction products were used as template in each reaction. The fold changes in target gene expression were normalized using Actin II (Xu et al., 2018). All primers used in this study are listed in Supplementary table. Five to seven biological replicates were included in assays.

RNA-seq service was provided by oebiotech company (www.oebiotech.com). Total RNA of three biological replicates of WT mock, inoculated WT leaf samples, irWRKY3 mock, inoculated irWRKY3 leaf samples were isolated with TRIzol reagent.

Sequencing was performed at 8 G depth and mapped to the *N. attenuata* reference genome sequence. The relative abundance of the transcripts was measured with the FPKM (RPKM) method. The differential expression between WT and irWRKY3 with or without *A. alternata* inoculation samples with a cutoff of 2-fold change.

### ChIP-qPCR assay

ChIP assays were performed using an EpiQuik Plant ChIP Kit (www.epigentek.com). Briefly, 7-week-old seedlings over-expressing *NaWRKY3-eGFP* were harvested with *A. alternata* at 1 dpi. Chromatin was isolated from 3 g leaf samples and sonicated with a Bioruptor pico-diagenode for 90 min. After coating with anti-GFP (1:1000, Abcam ab290) antibodies, NaWRKY3-eGFP protein/DNA complexes were immunoprecipitated according to the manufacturer’s instructions. The enriched DNA fragments were detected by qPCR using the specific primers listed in Supplementary table. The Actin II promoter was used as a negative control. All ChIP assays were performed with three biological replicates.

### Electrophoretic mobility shift assay (EMSA) assays

EMSA experiments were performed using an EMSA/Gel-Shift Kit (Beyotime) according to the manufacturer’s instructions. The NaWRKY3 contained two WRKY domains were cloned into the pET-28a (containing six His tag) vectors. The recombinant His-NaWRKY3 proteins were induced by 3 mM IPTG and expressed in *Escherichia coli* strain BL21. These recombinant proteins were purified with Ni-NTA agarose. Oligonucleotide probes were labeled by 5’-end biotins, and competitors were unlabeled in the binding assays (sequences are listed in Supplementary table). Equal amounts of His-NaWRKY3 proteins (1 µL) were incubated according to the manufacturer’s instructions in PCR volumes for 20 min at room temperature. The DNA-protein complexes were separated and transferred to a nylon membrane (Thermo Fisher Scientific). After UV cross linking, the biotin signal was detected according to the manufacturer’s instructions.

### Dual-LUC assays

Dual-LUC assays were performed in *N. benthamiana* leaves (Hellens et al., 2005) with pCAMBIA3301-LUC vector system which contained a *Renilla luciferase* (*REN*) gene driven by the 35S promoter as an internal control and a *firefly luciferase* (*LUC*) gene driven by the promoter of target gene. The target gene promoter fragments (the linear of fragment range from 1.1 kb to 3 kb from ATG site) were cloned and infused into pCAMBIA3301-LUC through the *Pst I* and *Nco I* sites, creating NaACS1_pro_::LUC, NaACO1_pro_::LUC, NaLOX3_pro_::LUC, NaF6’H1_pro_::LUC, NaBBL28_pro_::LUC, NaRboh D_pro_::LUC, L2_pro_::LUC vectors, independently. *Agrobacterium* carrying each reporter together with 35S::NaWRKY3-eGFP or 35S::eGFP were co-transformed into *N. benthamiana* leaves. The LUC activity of plant extracts was analyzed using a microplate reader (Tecan infinite M200 PRO) with commercial LUC reaction reagents following the manufacturer’s instructions (Yeasen Biotechnology). LUC activity was normalized with Renilla luciferase activity (LUC/REN). At least three replicates were performed for each experiment.

## Accession Numbers

Sequence data from this article can be found in the GeneBank under accession numbers: *NaWRKY3* (XM_019394677.1), *NaACS1* (AY426752), *NaLOX3* (AY254349), *NaACO1* (AY426756), *NaF6’H1* (KF771989), *NaBBL28* (XM_019378340.1), lncRNA *L2* (XR_002068323.1), *NaRboh D* (XM_019378071.1).

## Supplemental Data

**Supplementary table**

Primers and probes used in this study.

**Supplemental Fig. 1** The expression of *NaAOS* after *A. alternata* inoculation in WT and irWRKY3 plants, and generation of *NaWRKY3-eGFP* over-expression plants.

**Supplemental Fig. 2** The expression of lncRNAs after *A. alternata* inoculation in WT and irWRKY3 plants.

**Supplemental Fig. 3** lncRNA *L2* is required for *N. attenuata* resistance against *A. alternata.*

**Supplemental Fig. 4** *NaBBL28* is required for *N. attenuata* resistance against *A. alternata.*

**Supplemental Fig. 5** Silencing *NaBBL28* decreased HGL-DTGs.

**Supplemental Fig. 6** DAB staining of *A. alternata*-infected WT and irRboh D plants.

**Supplemental Fig. 7** Alignment of the *Arabidopsis thaliana* AtWRKY33 and *N. attenuata* NaWRKY3 protein sequences.

## Acknowledgments and Funding

We thank Prof. Ian T. Baldwin (Max-Planck Institute for Chemical Ecology, Jena, Germany) for providing irWRKY3 and irRboh D transgenic *N. attenuata* seeds and Biological Technology Open Platform of Kunming Institute of Botany for greenhouse and instrument services. This project was supported by the National Natural Science Foundation of China (NSFC Grant No. 31670262) and top-talent recruitment program of Yunnan to Prof. Jinsong Wu.

## Author Contributions

Jinsong Wu conceived the project; Jinsong Wu and Zhen Xu designed the experiments and prepared the manuscript. Zhen Xu performed most of experiments and analyzed the data. Shuting Zhang assisted in experiments of JA and ethylene parts and discussed the results.

## Conflict of Interest

All authors declare that they have no conflict of interest.

